# Long-term engraftment of adult hematopoietic progenitors in a novel model of humanized mice

**DOI:** 10.1101/2023.10.02.560534

**Authors:** Chun I Yu, Rick Maser, Florentina Marches, Jacques Banchereau, Karolina Palucka

## Abstract

Pre-clinical use of humanized mice transplanted with CD34^+^ hematopoietic progenitor cells (HPCs) is limited by insufficient engraftment with adult HPCs. Here, we developed a novel immunodeficient mice based in NOD-SCID-*Il2γc*^-/-^ (NSG) mice to support long-term engraftment with human adult HPCs and tissue colonization with human myeloid cells. As both Flt3L and IL-6 are critical for many aspects of hematopoiesis, we knock-out mouse *Flt3* and knock-in human *IL6* gene. The resulting mice showed an increase in the availability of mouse Flt3L to human cells, and a dose-dependent production of human IL-6 upon activation. Upon transplantation with low number of human HPCs from adult bone marrow, these humanized mice demonstrated a significantly higher engraftment with multilineage differentiation of human lymphoid and myeloid cells. Furthermore, higher frequencies of human lymphoid and myeloid cells were detected in tissues at one year after adult HPC transplant. Thus, these mice enable studies of human hematopoiesis and tissue colonization over time.

**Summary:** Pre-clinical use of humanized mice is limited by insufficient engraftment with adult hematopoietic progenitor cells (HPCs). Here, we developed a novel immunodeficient mice which support long-term engraftment with adult bone marrow HPCs and facilitate building autologous models for immuno-oncology studies.

## Introduction

Realizing the clinical promise of cancer immunotherapy is hindered by gaps in our knowledge of *in vivo* mechanisms underlying treatment response as well as treatment limiting toxicity. Preclinical *in vivo* model systems and technologies are required to address these knowledge gaps and to surmount the challenges faced in the clinical application of immunotherapy. Mice are commonly used for basic and translational research to support development and testing of immune interventions, including for cancer. Yet, despite some remarkable advances, current mouse models share an obvious and important limitation when it comes to translational potential: their immune system is that of a mouse. To overcome this, mice can be humanized through genetic approaches leading to expression of human proteins, or through cellular approaches based on transplantation of human cells such as peripheral blood mononuclear cells (PBMCs), fetal bone marrow/liver/thymus tissues (BLT), or human CD34^+^ hematopoietic progenitor cells (HPCs) (Garcia, 2016; Matsumura et al., 2003; Traggiai et al., 2004). However, the allogenic context between human immune cells and cancer cells imposes a confounding influence on the anti-tumor effect. To overcome this, an autologous humanized tumor model is needed, where mice are transplanted with bone marrow derived HPCs followed by implantation of patient-derived xenograft (PDX) tumor from the same patient (Chiorazzi et al., 2023; Fu et al., 2017).

Humanization by transplanting fetal human CD34^+^ HPCs into immunodeficient mice bearing the *Il2rg* mutation has demonstrated an human engraftment including T cells (Matsumura et al., 2003; Traggiai et al., 2004) and it has since contributed significantly to human immunology research (Ito et al., 2018; Legrand et al., 2009; Saito et al., 2020; Theocharides et al., 2016). However, model based on NOD-SCID-*Il2γc*^-/-^ (NSG) cannot efficiently support the generation of a diverse and fully functional human immune system for efficient preclinical and translational immunology studies. Various efforts have been made on genetically engineering to express human cytokines to cross mouse and human barriers (Willinger et al., 2011a). Human growth factors including Granulocyte-Colony Stimulating Factor (G-CSF), Granulocyte Macrophage-Colony Stimulating Factor (GM-CSF), Macrophage-Colony Stimulating Factor (M-CSF), Interleukin (IL)-3, IL-6, IL-15, IL-34 and Stem Cell Factor (SCF) have been shown to impact human hematopoiesis which led to adequate development of human hematopoietic lineages in humanized mice (Cosgun et al., 2014; Hanazawa et al., 2018; Huntington et al., 2011; Jangalwe et al., 2016; Mathews et al., 2019; Rathinam et al., 2011; Rongvaux et al., 2014; Theocharides et al., 2016; Willinger et al., 2011b; Yu et al., 2017). Nevertheless, few humanized mouse models have reported of human engraftment by adult HPCs and mature monocyte development (Chiorazzi et al., 2023; Saito et al., 2016). A general lack of multilineage human engraftment was reported in mice transplanted with limited number of adult HPCs (Beyer and Muench, 2017).

Flt3L and IL-6 are cytokines important for hematopoiesis. Loss-of-function mutation in murine *Flt3* leads to a decrease in mouse dendritic cells (DCs) and an accumulation of mouse ligand in the serum (Karsunky et al., 2003; McKenna et al., 2000; Sitnicka et al., 2002). This could lead to the creation of “space” in peripheral tissues enabling colonization with human cells upon HPC transplant as well as increased availability of murine Flt3L to the human receptor, supporting development of human immune cells including DCs (Maraskovsky et al., 2000) and B cells (Miller et al., 1999). IL-6 lacks cross-reactivity between mouse and human and plays key roles in the innate and adaptive immune systems, including HPC survival (Moldenhauer et al., 2008), monocyte differentiation to macrophages by facilitation of autocrine CSF-1 internalization (Chomarat et al., 2000), and the differentiation of follicular helper T cells, and antibody producing plasma cells (Das et al., 2016; Jego et al., 2003; Nurieva et al., 2009; Papillion et al., 2019). To overcome the limitation on the engraftment with adult HPCs, we knocked-out (KO) mouse *Flt3* in order to increase the availability of mouse Flt3L (which can act via human receptor) to human cells, thereby improving the long-term development of human myeloid cells upon transplantation with human CD34^+^ HPCs (Li et al., 2016). Furthermore, we genetically engineered NSG mice to express human IL-6. This novel mouse model yielded a higher human engraftment and support long-term multilineage development of human immune cells when transplant with adult human CD34^+^ HPCs.

## Results and Discussion

### Construction of murine *Flt3* KO and human *IL6* KI mice via CRISPR

We first used the CRISPR/Cas9 system to generate a NSG mice with *Flt3* KO (NSGF) (Wang et al., 2013). Founder mice carrying a chromosomal deletion at the exon 3 were backcross to NSG mice and inbred to obtain homozygous *Flt3*^*-/-*^ allele confirmed by PCR and Sanger sequence (**Fig. 1 A; and Fig. S1 A**). To verify the impact of *Flt3* KO on mouse DCs, we analyzed different subsets of mouse DCs including CD317^+^ plasmacytoid DCs (pDCs), CD11c^+^ conventional DCs (cDCs). As expected, we observed an 80% decrease in both pDCs and cDC in the bone marrow, spleen, and lungs of NSGF mice in comparison to age and gender matched NSG mice by FACS (**Fig. 1 B; and Fig. S1, B and C**). Consequently, we also observed an increase amount of mouse Flt3L in the plasma of NSGF mice (**Fig. 1 C**).

**Figure 1.**
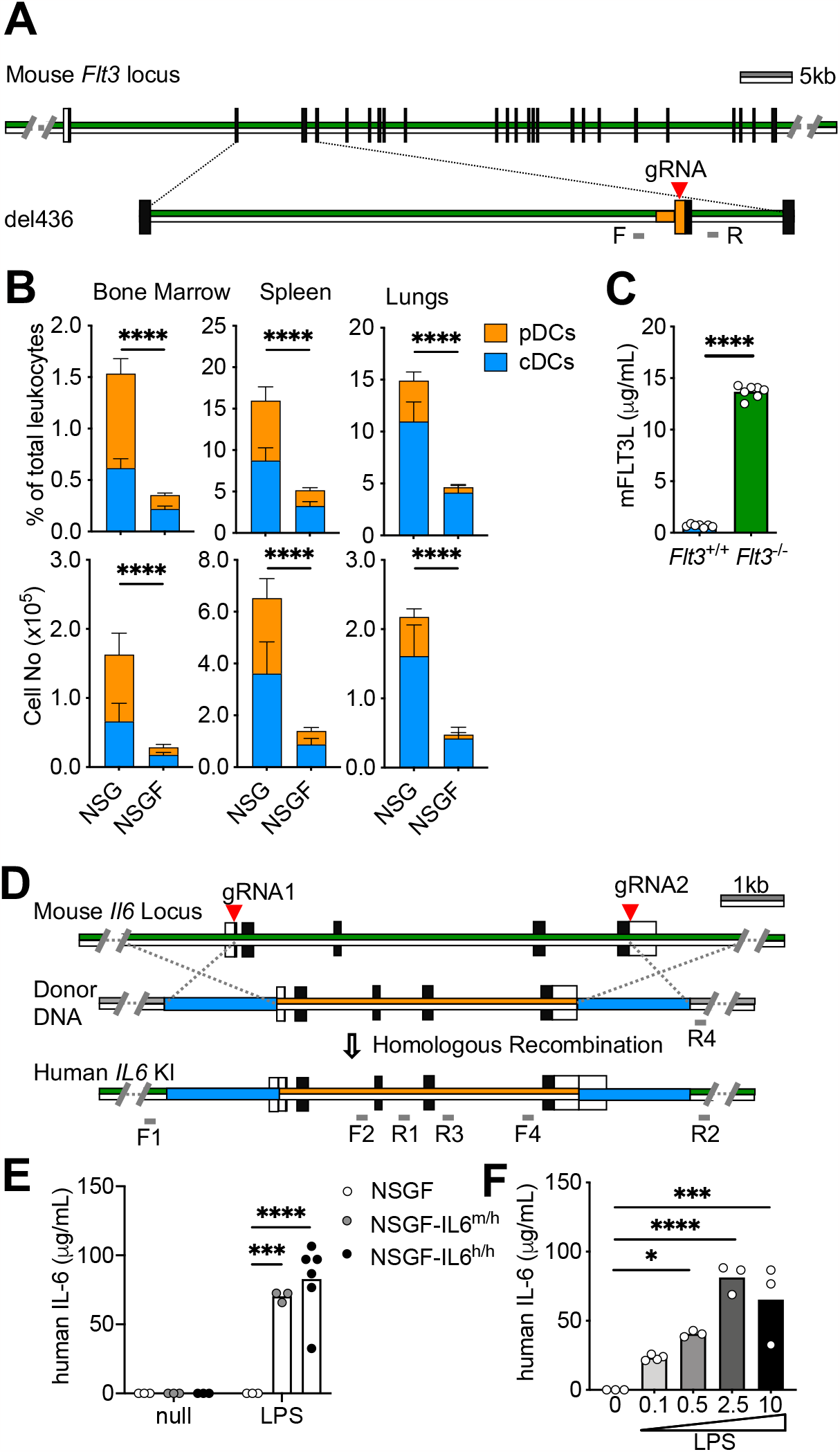
Genetic engineering of novel NSG mice via CRISPR. **(A)** Schematic representation of CRISPR/Cas9 at mouse *Flt3* locus and the chromosomal deletion at the exon 3. Green lines represent the mouse sequences and orange lines represent the mutated sequences. Black boxes represent the coding region and white box represent the non-coding region. gRNAs are labeled with red arrowhead and the primer pairs F/R are in grey. **(B)** The summary of percentage (upper) and total number (lower) of mouse pDCs and cDCs from bone marrow, spleen and lungs of 8-10 wk of mice (mean±SD, n=7). **(C)** Mouse Flt3L in the plasma of 8-10 wk of mice were analyzed by ELISA (mean, n=7). **(D)** Schematic representation of CRISPR/Cas9 at mouse *Il6* locus, the vector for human *IL6* donor DNA and targeted allele with homologous recombination. Green lines represent the mouse sequences and orange lines represent the human *IL6* sequences. Black boxes represent the coding region and white box represent the non-coding region. gRNAs are labeled with red arrowhead. Blue boxes represent the left and right homologous arms. One of the primer pairs F1/R1 and F2/R2 in grey are located outside of the homologous arms, respectively. **(E)** Human IL-6 production (mean, n=3-6) in the plasma of mice treated with 10 μg of LPS i.p. for 2 hours. **(F)** Human IL-6 production (mean, n=3) in the plasma of mice treated with different dose of LPS (0.1, 0.5, 2.5, 10 μg) i.p. for 2 hours.

Next, we generated human *IL6* knockin (KI) in replace of mouse ortholog using CRISPR/Cas9 gene targeting in zygotes and inserting the full-length of human *IL6* gene sequence into exon 1 and exon 5 of mouse *Il6* locus via homologous recombination (**Fig. 1 D**). Founder mice were selected first with PCR assays targeting 5’ and 3’ junctions and full-length sequence between two homology arms. In addition, we also confirmed the absence of plasmid donor sequences to discern correct on-target single copy integration events from random or multi-copy targeting events. Sequencing of these PCR products confirmed proper targeting of human *IL6*. Five founder mice with on-target single copy integration events of human *IL6* KI were identified (**Fig. S1 D**). Founder mice with human *IL6* KI were first bred to NSGF mice and then intercrossed to generate homozygous animals for *IL6* targeted mutation (NSGF6). Human IL-6 was found in the plasma of mice with *IL6*^m/h^ and *IL6*^h/h^ genotype but not wildtype mice upon i.p. challenge with LPS in a dose-dependent manner (**Fig. 1, E and F**).

### Human *IL6* KI boost HPC engraftment and human monocyte differentiation

As IL-6 promote HPC survival (Moldenhauer et al., 2008), we first analyzed engraftment with cord blood CD34^+^ HPCs to test whether human *IL6* KI improves humanization. While comparable engraftment was found in the blood of mice transplanted with high number of HPCs (1x10^5^) from cord blood, humanized (h)NSGF6 mice but not hNSG mice were consistently engrafted when transplanted with low number of HPCs (1x10^4^) from cord blood (**Fig. 2 A**). The difference in engraftment was found in the blood over a six-month follow up (**Fig. 2 B**). Furthermore, the improvement was found in both CD33^+^ myeloid cell and CD3^+^ T cell development with a significant increase in the absolute cell counts (**Fig. 2 C**). In addition, we also observed the presence of different blood monocyte subsets with a significant increase in CD16^+^ non-classical monocytes (**Fig. 2 D**), and predominately CD45RA^+^ CCR7^+^ naive T cells were found in the blood at 16-week after HPC transplant (**Fig. 2 E**). Thus, we conclude that mouse *Flt3* KO and human *IL6* KI improves the engraftment of human immune cells both quantitatively and qualitatively upon transplant with human cord blood derived CD34^+^ HPCs.

**Figure. 2.**
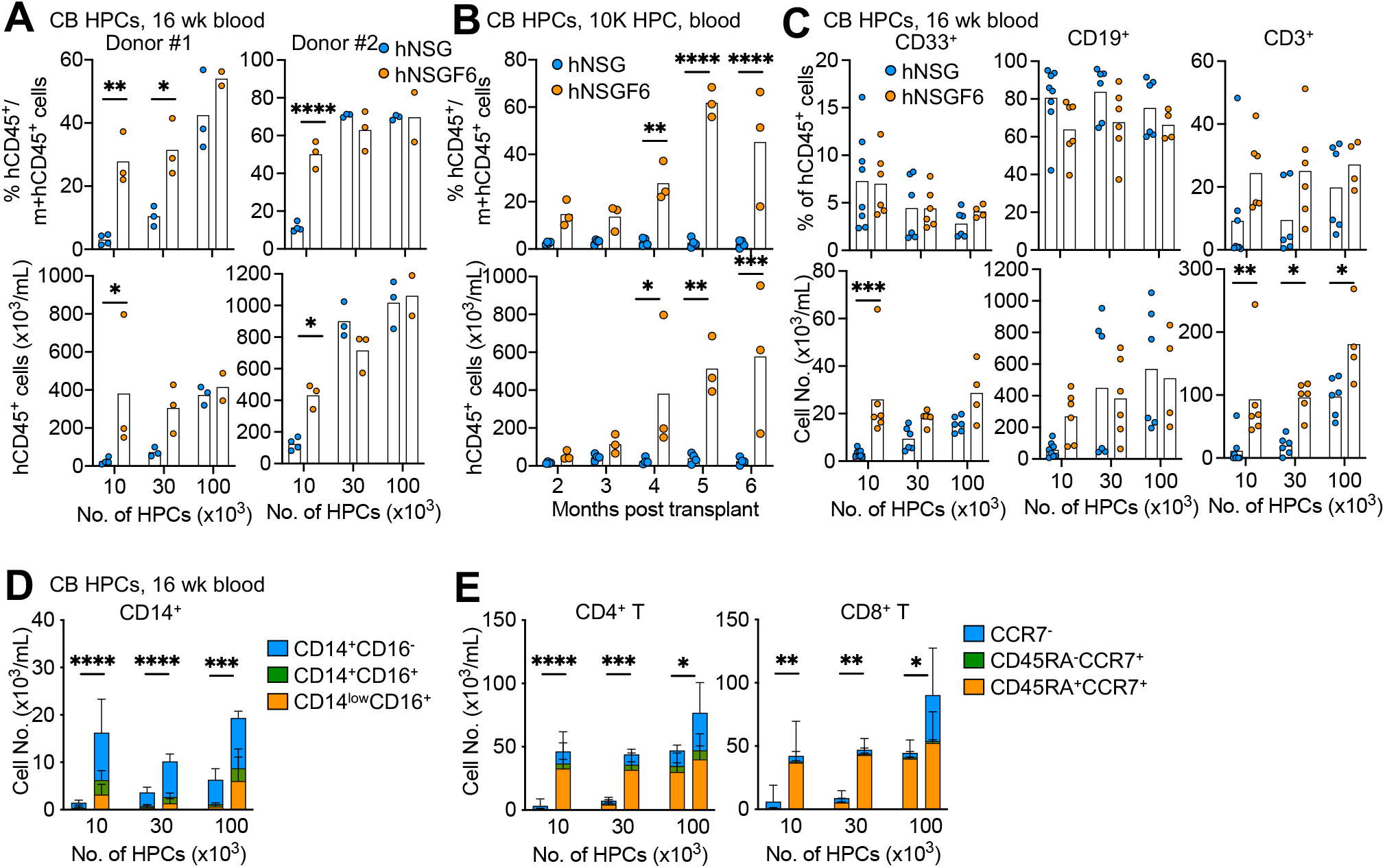
Improved human engraftment by HPCs from cord blood. **(A)** NSG or NSGF6 mice were transplanted with 1x10^4^, 3x10^4^, 1x10^5^ HPCs from cord blood at 4-week of age. Human engraftment in the blood by the mean percentage (upper) and absolute number (lower) of hCD45^+^ cells at 16 weeks after transplant. n=2-4 mice from two independent experiments. 2way ANOVA. **(B)** The kinetics of mean human engraftment in the blood in hNSG or hNSGF6 mice after transplant of 1x10^4^ cord blood HPCs. n=3-4 mice. 2way ANOVA. **(C)** The mean percentage (upper) and absolute number (lower) of human CD33^+^ myeloid cells, CD19^+^ B cells, and CD3^+^ T cells in the blood at 16 weeks after transplant. n=4-8 mice pooled from two donors. 2way ANOVA. **(D)** The absolute number of human CD14^+^ monocyte subsets in the blood at 16 weeks after transplant (mean±SD, n=4-8). 2way ANOVA. **(E)** The absolute number of human CD4^+^ CD3^+^ (left) or CD8^+^ CD3^+^ T cell subsets in the blood at 16 weeks after transplant (mean±SD, n=4-8). 2way ANOVA.

As the three subsets of human monocytes generated through distinct pathways with diverse function: CD14^+^CD16^-^ classical, CD14^+^CD16^+^ intermediate, and CD14^low^CD16^+^ non-classical monocytes (Guilliams et al., 2018), we want to test whether human *IL6* KI are indeed essential for monocyte differentiation. To this end, we set up an experiment to compare NSGF6 mice and NSGF mice using high number of cord blood HPCs (1x10^5^). As expected, similar engraftment with different lineages of human immune cells were found in various tissues of hNSGF and hNSGF6 mice with slight variability found among individual mouse (**Fig. 3 A**). However, in comparison to hNSGF mice, significantly higher frequencies of CD14^+^ cells were found in the spleen and lungs of hNSGF6 mice (**Fig. 3 B**). We also found higher frequencies of both CD14^+^CD16^+^, and CD14^low^CD16^+^ monocytes in the spleen and lungs of hNSGF6 mice engrafted with cord blood HPCs (**Fig. 3 C and D**). Thus, our data collaborated with others (Hanazawa et al., 2018) and demonstrate that human IL-6 is important for the development of CD16^+^ monocytes.

**Figure. 3.**
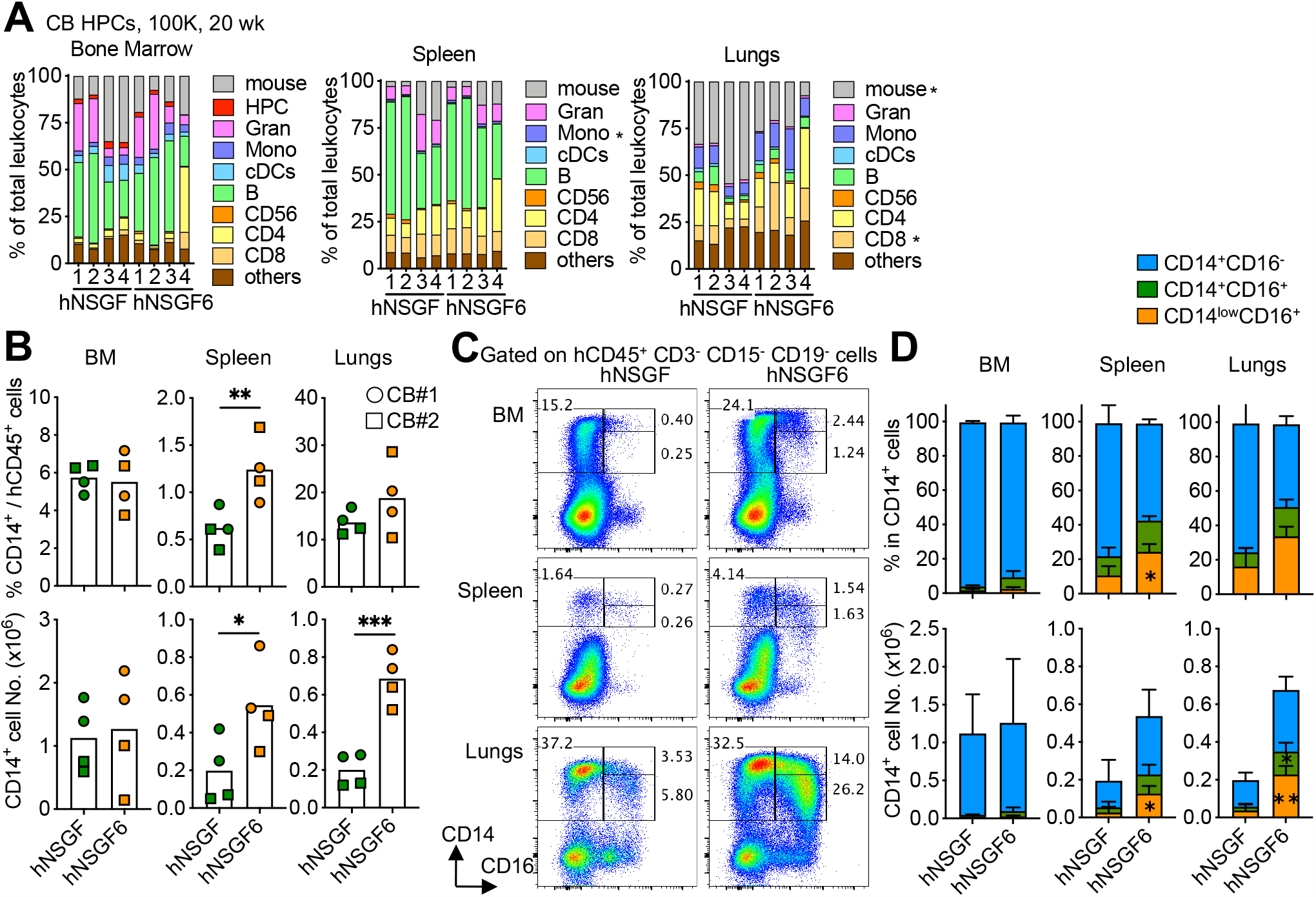
Human IL-6 supports human monocyte differentiation. **(A)** The percentage of human immune cell type in total CD45^+^ leukocytes (mouse and human) in the bone marrow, spleen and lungs of hNSGF and hNSGF6 at 20-week post-transplant with cord blood HPCs. n=4 from two donors. Two-tailed t-test. **(B)** Summary of the mean percentage (upper panels) and absolute number (lower panels) of CD14^+^ cells in mice. n=4 from two donors. Two-tailed t-test. **(C)** Representative FACS plots of human monocyte subsets. **(D)** Summary of CD14^+^ cell subsets (mean±SD, n=4). 2way ANOVA.

### Murine *Flt3* KO and human *IL6* KI support long-term engraftment by adult HPCs

The key question was if our novel model enables engraftment of adult bone marrow HPCs which thus far has proven difficult in other models. To this end, we transplanted 1x10^5^ bone marrow derived CD34^+^ HPCs into sublethally irradiated NSG, and NSGF6 mice. As shown in **Fig. 4**, we found a significant higher hCD45^+^ engraftment in the blood of hNSGF6 mice than hNSG mice (**Fig. 4 A; and Fig. S2**). Human CD33^+^ myeloid cells, CD19^+^ B cells, and CD3^+^ T cells were all present in significant higher numbers in the blood of hNSGF6 mice than hNSG mice (**Fig. 4 B**). Furthermore, hNSGF6 mice demonstrated superior overall engraftment than hNSG mice over one-year follow up (**Fig. 5 A**). hCD33^+^ myeloid cells were consistently present in the blood of hNSGF6 mice (**Fig. 5 B**). In contrast to the persistent lack of T cell developments in hNSG mice, circulating CD3^+^ T cells were found hNSGF6 as early as 16 weeks, reached plateau at 24 weeks and remained persistent up to one year follow-up (**Fig. 5 C and C**). Additionally, when we looked into the T cell phenotype, we found a stable presence of CD45RA^+^CCR7^+^ naïve T cells despite an increase in expansion or accumulation of CD4^+^ T cells with memory phenotype (**Fig. 5 D**). Finally, we analyzed the human immune composition in the tissues of humanized mice at one year after engraftment with bone marrow HPCs. In comparison to hNSG mice, hNSGF6 mice showed significantly higher percentage of CD11c^+^ cDCs and CD56^+^ NK cells in bone marrow; and drastic increase in many lineages of human immune cells including CD15^+^ or CD66b^+^ granulocytes, CD14^+^ monocytes, CD11c^+^ cDCs, CD19^+^ B cells, CD56^+^ NK cells and CD4^+^ T cells in the spleen or lungs (**Fig. 5 E and Fig. S 3**). Consistently, we also found a significant higher frequency of total CD14^+^ monocytes with CD14^+^CD16^+^, and CD14^low^CD16^+^ phenotype in the spleen and lungs of hNSGF6 mice (**Fig. 5 F**). Thus, human *IL6* KI combined with *Flt3* KO improve the engraftment and long-term development of human immune cells upon transplant with adult human CD34^+^ HPCs.

**Figure. 4.**
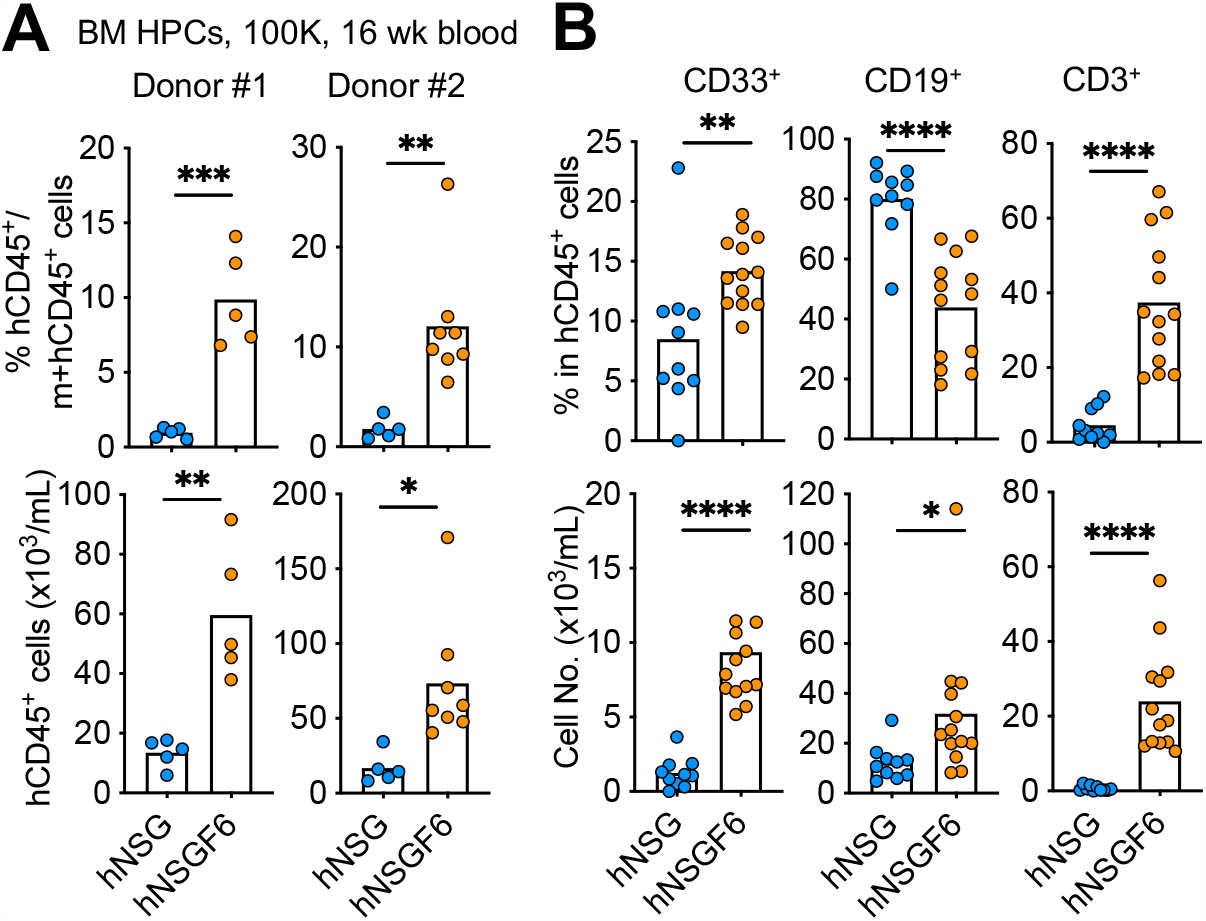
Improved human engraftment by adult HPCs. **(A)** Humanized mice were generated by engrafting NSG and NSGF6 mice with 1x10^5^ bone marrow HPCs. The mean percentage (upper) and absolute number (lower) of hCD45^+^ cells at 16 weeks after transplant. n=5-8 mice from two donors. Two-tailed t test. **(B)** The mean percentage (upper) and absolute number (lower) of human CD33^+^ myeloid cells, CD19^+^ B cells and CD3^+^ T cells in the blood at 16 weeks after transplant. n=10-13 mice pooled from two donors. Two-tailed t test.

**Figure. 5.**
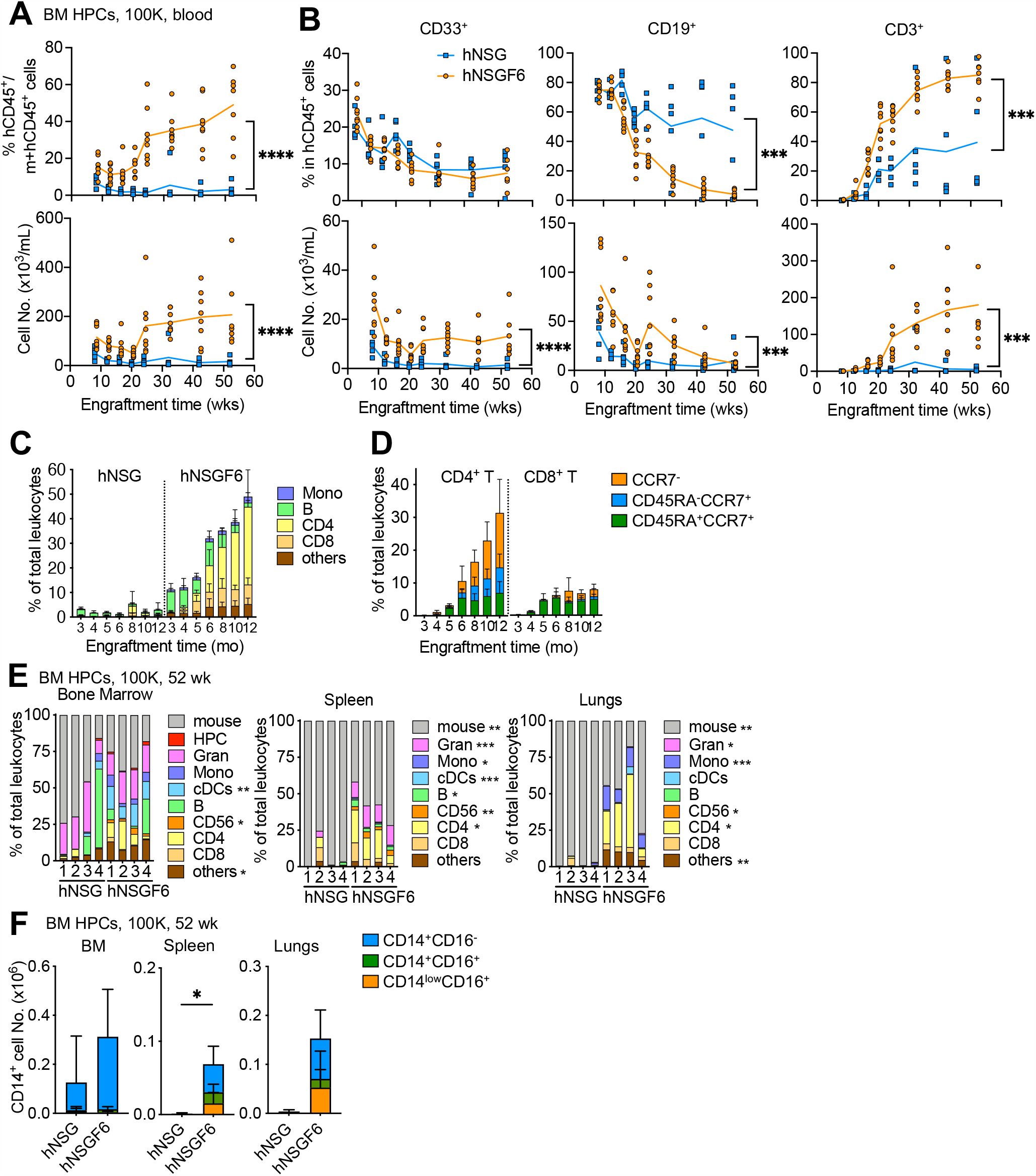
Human *IL6* KI support long-term engraftment by adult HPCs. **(A)** The kinetics of mean percentage (upper panel) and absolute number (lower panel) of hCD45^+^ cells in the blood of NSG and NSGF6 mice at different time points after transplant with 1x10^5^ bone marrow HPCs. n=5-8 from one donor. Two-way ANOVA. **(B)** The kinetics of mean percentage (upper panel) and absolute number (lower panel) of human CD33^+^ myeloid cells, CD3^+^ T cells, and CD19^+^ B cells (mean, n=5-8 mice) in the blood. One-way ANOVA. **(C)** The percentage of different human immune cell types in total CD45^+^ leukocytes (mouse and human) analyzed at different time points after HPC transplant in the blood (mean±SD, n=5-8 mice). **(D)** Kinetics of human CCR7^+^ CD45RA^+^ naïve, CCR7^+^ CD45RA^-^ central memory, and CCR7^-^ effector memory CD4^+^ or CD8^+^ T cell frequency in the blood of hNSGF6 mice (mean±SD, n=5-8 mice). **(E)** The percentage of different human immune cell types in total CD45^+^ leukocytes (mouse and human) in the bone marrow, spleen and lungs analyzed at one year post transplant by FACS. n=4 mice from one bone marrow donor. Two-tailed t-test. **(F)** Summary of the CD14^+^ cell subsets (mean±SD, n=4 mice). Two-tailed t-test.

In summary, we have developed a novel strain of humanized mouse model which enables multilineage development of human immune cells in tissues when engrafted with adult HPCs. Hence, our new model should improve generation of autologous models and enhance our ability to investigate human tumor-human immune system interactions in humanized mice.

## Materials and methods

### Antibodies and reagents

Antibodies were obtained from BD (Franklin Lakes, NJ), Biolegend (San Diego, CA), or ThermoFisher (Waltham, MA). Detail information on biologicals and reagents was listed in **Table S1**.

### Genetic engineering of mouse models

All mice were obtained from The Jackson Laboratory (Bar Harbor, ME). The protocol on genetic engineering of mouse models was reviewed and approved by the Institutional Animal Care and Use Committee at The Jackson Laboratory (14005). Mouse *Flt3* KO mice (NOD.Cg-Prkdc^scid^ *Il2rg*^tm1Wjl^-*Flt3*^em1Akp^; NSGF ; RRID:IMSR JAX:035842) were generated by CRISPR using Cas9 mRNA and sgRNAs (5’-AAGTGCAGCTCGCCACCCCA-3’) targeting exon 3 of mouse *Flt3* in fertilized eggs of NSG mice (NOD.Cg-Prkdc^scid^ *Il2rg*^tm1Wjl^/SzJ; RRID:IMSR JAX:005557). The blastocysts derived from the injected embryos were transplanted into foster mothers and newborn pups were obtained. Founders and F1 littermates were tail tipping and tested for successful gene-knockout by PCR and Sanger sequencing. Mice carrying a null deletion were backcrossed to NSG for two generations and were then interbred until all offspring were homozygous for *Flt3* KO mutation. Human *IL6* KI mice (NOD.Cg-*Prkdc*^*scid*^ *Il2rg*^*tm1Wjl*^*-Flt3*^*em1Akp*^*Il6*^*emX(IL6)Akp*^; NSGF6) were generated using CRISPR/cas9 system. sgRNAs were designed to target exon 1 (5’-TGCAGAGAGGAACTTCATAG-3’ or 5’-AGGAACTTCATAGCGGTTTC-3’) *and* exon 5 (5’-ATGCTTAGGCATAACGCACT-3’) of mouse *Il6* locus. Cas9 mRNA, sgRNAs targeting mouse *Il6* locus and recombinant human *IL6* DNA were coinjected into fertilized NSGF oocytes. Human *IL6* was inserted into exon 1 and exon 5 via homologous recombination. The resulting founders, carrying human *IL6* were bred to NSGF mice for two generations, and were then interbred until all offspring were homozygous for *Il6* targeted mutation. For human IL-6 production, mice at 6-8 week of age were treated with 0.1-10 μg of LPS (InvivoGen, San Diego, CA) i.p. and euthanized after 2 hours for plasma.

### Genotyping assays

For all founders, the mice were typed by PCR reaction for the presence or absence of the *Flt3* KO and human *IL6* KI sequence using primers listed in **Table S2**. Genomic DNA was isolated from tail biopsies using the NucleoSpin Tissue kit (TaKaRa Bio USA, San Jose, CA) following manufacture protocol. All primers were purchased from Eurofins Genomics (Louisville, KY). PCR reactions were performed with PrimSTAR GXL DNA polymerase (TaKaRa Bio USA) with 1 μl of genomic DNA and 0.2 μM primers. A common PCR thermal-cycle amplification program was used for all primer pairs (10 sec at 98°C and 1-3 min at 68°C for 30 cycles). PCR products were run on agarose gels and size was estimated with comparison to a DNA mass ladder. In some experiment, the remaining PCR products were extracted with PCR cleanup kit (TaKaRa Bio USA) and sequenced to confirm the correct gene sequence.

### Humanized mice

Humanized mice were generated on mice obtained from The Jackson Laboratory (Bar Harbor, ME). All protocols were reviewed and approved by the Institutional Animal Care and Use Committee at The Jackson Laboratory (14005) and University of Connecticut Health Center (101163-0220 and 102195-1122; Farmington, CT). Female mice were sub-lethally irradiated (10 cGy per gram of body weight) using gamma irradiation at four weeks of age. Human CD34^+^ HPCs isolated from full-term cord blood (Lonza) or adult bone marrow (Lonza) were given by tail-vein intravenous (i.v.) injection in 200 μL of PBS with 1 μg/mL of anti-CD3 antibody (OKT3, Biolegend). Monthly post HPC transplant, one capillary tube of blood was collected with heparin from the mice to evaluate blood engraftment and mice were euthanized according to the individual experimental design.

### Tissue processing for flow cytometry

For immunophenotype, tissues including blood, bone marrow, spleen and lungs were collected. Bone marrow was harvested from femur and tibia by flushing out the marrow. Spleen was digested with 50 μg/ml of Liberase (Millipore Sigma) and DNase I (Millipore Sigma) for 10 min at 37°C. Lungs were digested with 50 μg/ml of Liberase and DNase I for 30 min at 37°C, followed by mechanical dissociation with GentleMACS (Miltenyi Biotec, Bergisch Gladbach, Germany). Single cell suspensions were made, and the debris was removed by filtering through 70 μm cell strainers. Cells were first treated with RBC lysis buffer (Biolegend) to remove red blood cells. Total cell counts were measured in hemocytometer with 0.4% trypan blue. Single cell suspensions were treated with Fc blocker (BD), and then stained on ice with antibody cocktail for 30 minutes. After washing twice with PBS, samples were resuspended in buffer (PBS, 2% FBS, 2 mM EDTA), acquired on a Symphony A5 (BD) and analyzed with FlowJo software (BD).

For the analysis of mouse DCs, cells were stained with antibodies to mouse CD45-BV650 (30-F11, BD), CD3-PE-CF579 (145-2C11, BD), CD19-PE-CF579 (ID3, BD), CD103-PerCP-Cy5.5 (M290, BD), F4/80-PE-Cy7 (BM8, Biolegend), Ly-6G/Ly-6C-Pacific Orange (RB6-8C5, ThermoFisher), MHC class II-FITC (10-3.6, BD), CD11c-V450 (HL3, BD), CD8-PE (53-6.72, BD), and CD317-APC (927, Biolegend). For human engraftment in the blood, cells were stained with mCD45-BV650 (30-F11, BD), and antibodies to human CD45-BV510 (HI30, BD), CD14-AF488 (HCD14, Biolegend), CD33-PE (P67.6, Biolegend), CD19-APC (HIB19, Biolegend) and CD3-APC-H7 (SK7, BD). Occasionally cells were stained with additional antibodies including CD45RA-PerCPCy5.5 (HI100, BD), CCR7-PE-Cy7 (3D12, BD), CD8-PB (RPA-T8, BD) and CD4-BUV395 (SK3, BD) for T cell phenotype. For cellular composition in tissues, cells were stained with antibody cocktail containing mCD45-BV650, hCD45-BV510, CD15-FITC (HI98, Biolegend), CD1c-PerCP-Cy5.5 (L161, Biolegend), CD33-PE, CD3-PE-CF594 (UCHT1, BD), CD19-PE-CF594 (HIB19, BD), CD14-PE-Cy7, CD141-APC (AD5-14H12, Miltenyi Biotec), CD66b-AF700 (G10F5, Biolegend), HLA-DR-APC-H7 (G46-6, BD), CD11c-V450 (B-ly6, BD), CD117-BV605 (104D2, Biolegend), CD11b-BV711 (D12, BD), CD303-BV785 (201A, Biolegend), and CD16-BUV395 (3G8, BD) in Brilliant Stain Buffer Plus (BD) for myeloid phenotype; and antibody cocktail containing antibodies to mCD45-BV650, hCD45-BV510, CD38-FITC (HB-7, Biolegend), CD45RA-PerCP-Cy5.5, CD138-PE (MI15, Biolegend), CD19-PE-CF594, CCR7-PE-Cy7, CD185-AF647 (RF8B2, BD), CD20-AF700 (2H7, Biolegend), CD8-APC-H7 (HIT8a, BD), CD279-BV421 (EH12.2H7, Biolegend), CD56-BV605 (NCAM16.2, BD), CD27-BV711 (M-T271, Biolegend), CD3-BV786 (SK7, BD) and CD4-BUV395 for lymphoid phenotype. Viability dye 7-AAD (Biolegend) were added to samples before acquisition on a flow cytometer.

### ELISA

ELSA were performed following manufacture protocol. For mouse Flt3L, plasma was tested with mouse Flt3L ELISA Duo Set from R&D systems. For human IL-6, plasma was tested with human IL-6 ELISA MAX Deluxe Set from Biolegend.

### Statistical analyses

Statistical analysis was performed in Prism (GraphPad, San Diego, CA). Legend is: ***P < 0.001, **P < 0.01, *P < 0.05, ns = not significant. Comparisons between any 2 groups were analyzed using the Mann-Whitney test or two-tailed t-test. Comparisons between any 3 or more groups were analyzed by analysis of variance (ANOVA).

## Supplemental material

Supplementary material including figures and tables. Figure S1 shows additional information on genetic engineering of novel NSG mice via CRISPR. Figure S2 shows evaluation of human engraftment in the blood of humanized mice by flow cytometry. Figure S3 shows evaluation of human myeloid compartment in the tissues of humanized mice by flow cytometry. Table S1 shows the list of biologicals and reagents used in this study. Table S2 shows the list of primers for mouse genotypes.

## Data availability statement

The data generated in this study are available within the article and its supplementary data files.

## Acknowledgements

We thank healthy donors for participation in our studies over the years. We thank Dr. Lenny Shultz for discussion; Pierre Authie, Patrick Metang, Vanessa KP Oliveria, Mayerlin Chalarca, Michael Michaud and PDX core at JAX-MG for help with mice; GET at JAX-MG for help with mouse model generation; Transgenic Genotyping Service at JAX-MG; Breeding Service and Research Animal Facility at JAX-MG; Molecular Diagnostics at JAX-MG; Comparative Medicine and Quality Service at JAX-MG; Comparative Medicine at UCHC. This work was supported by grants from The Jackson Laboratory Director Innovation Fund and NIH (NCI P30 CA034196 and R01 CA219880). CY, RM, JB and KP filed a patent on novel humanized mouse models via genetic editing of NSG mouse. The authors have no additional financial interests relevant to the work described in this study.

## Author contributions

CY designed and performed experiments, analyzed the data, and co-wrote the manuscript. RM designed and performed CRISPR experiments. FM constructed humanized mice. JB contributed to study design and manuscript writing. KP developed the concept, designed the study, analyzed the data and co-wrote the manuscript. All authors edited the manuscript.

## Abbreviations

BLT: fetal bone marrow/liver/thymus tissues
BM: bone marrow
CB: cord blood
cDCs: conventional DCs
Cas9: CRISPR-associated protein-9 nuclease
CRISPR: clustered regularly interspaced short palindromic repeats
DCs: dendritic cells
Flt3: FMS-like tyrosine kinase 3
Flt3L: FMS-like tyrosine kinase 3 Ligand
Gran: granulocytes
sgRNA: single-guide RNA
h: human or humanized
HPCs: hematopoietic progenitor cells
KI: knockin
m: mouse or murine
Mono: monocytes
NSG: NOD SCID gamma
NSGF: NSG mice with *Flt3* KO mutation
NSGF6: NSG mice with *Flt3* KO and human IL6 KI mutation
PBMCs: peripheral blood mononuclear cells
pDCs: plasmacytoid DCs
PDX: patient-derived xenograft

