## Supplemental figures and tables for "Long-term engraftment of adult hematopoietic progenitors in a novel model of humanized mice"

Figures S1 to S3
Tables S1 to S2

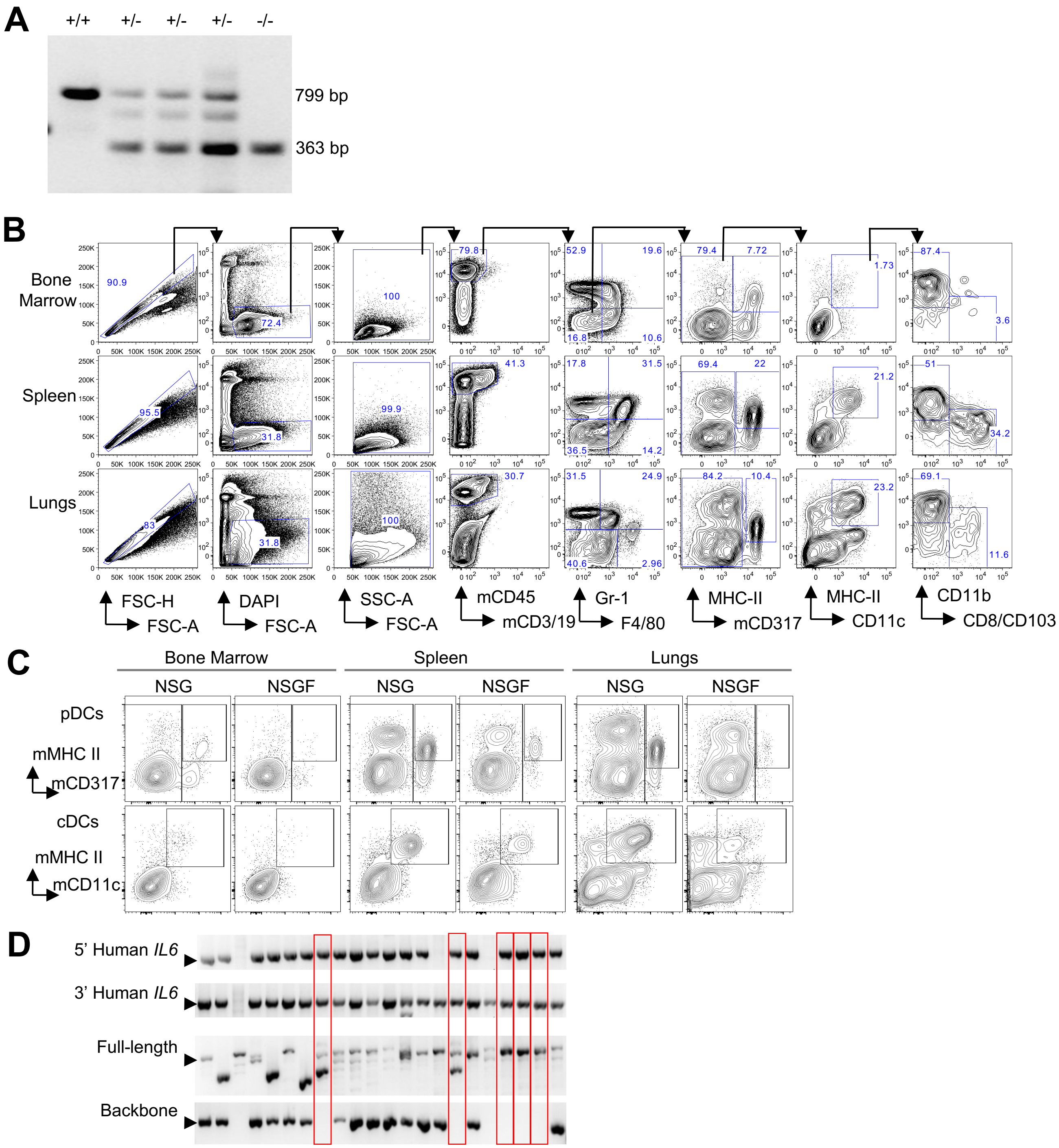


Figure S1. **Genetic engineering of novel NSG mice via CRISPR.**

**(A)** F1 littermates from mouse *Flt3* CRISPR KO were screened for *Flt3* mutant allele (363 bp) from wildtype allele (799 bp) by PCR.

**(B)** The gating strategy for mouse DCs on single cell suspension from bone marrow, spleen and lungs of NSG mice by flow cytometry. pDCs were gated as DAPI$^-$, mCD45$^+$, mCD3$^-$, mCD19$^-$, F4/80$^-$ and Gr-1$^-$ with expression of MHC class II and CD317. CD317$^-$ cells were further gated with MHC class II$^+$ and mCD11c$^{+}$ for cDCs. cDCs were subsequently divided into mCD8$^{+}$ or mCD103$^{+}$ cDC1 and mCD11b$^{+}$ cDC2.

**(C)** Representative FACS plots illustrate mouse pDCs and cDCs from bone marrow, spleen, and lungs of 8-10 wk NSG or NSGF mice (n=7).

**(D)** Founder mice from human *IL6* CRISPR KI were screened for mutant allele from wildtype allele by PCR. PCR products obtained with template of genomic DNA from tail tip. Potential founder mice indicated with red box were selected by positive PCR assay targeting 5' and 3' junctions and full length of human *IL6* KI sequence and negative for plasmid backbone.

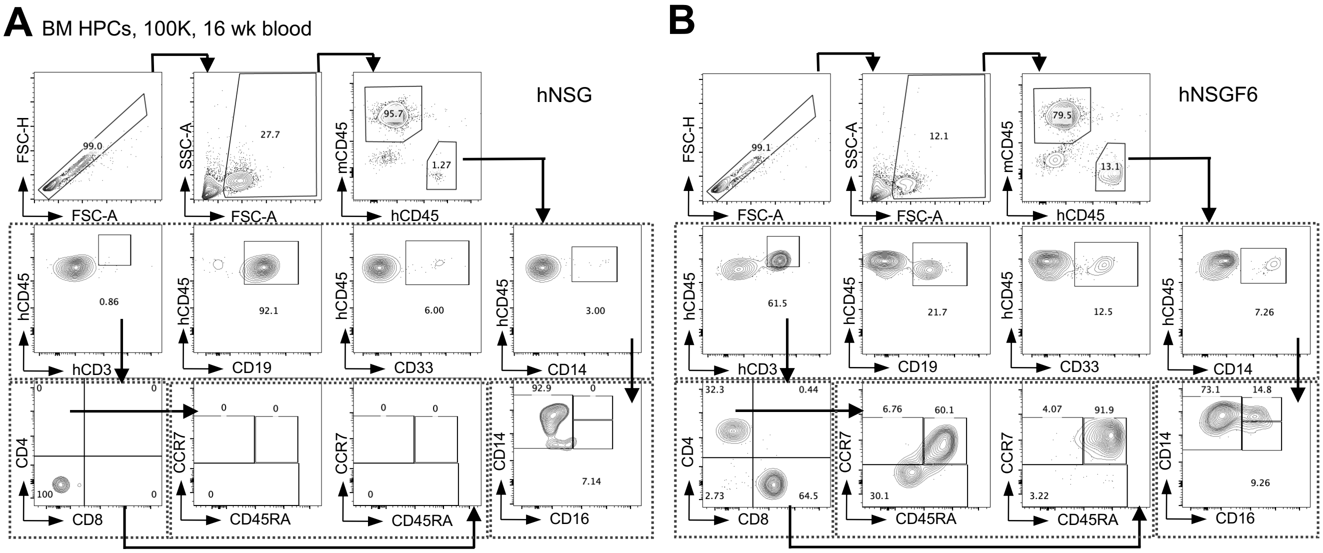


Figure S2. **Evaluation of the human engraftment in the blood of humanized mice by flow cytometry.** Humanized mice were generated by engrafting mice with $1x10^5$ bone marrow HPCs. Peripheral blood of mice was collected, stained with specific antibodies, and analyzed by FACS.

**(A)** The gating strategy for human engraftment in the blood of hNSG at 16 weeks after transplant. hCD45$^{+}$ cells were gated as mCD45$^{-}$ with expression of hCD45. hCD45$^{+}$ cells were further gated into CD14$^{+}$ monocytes, CD33$^{+}$ myeloid cells, CD19$^{+}$ B cells or CD3$^{+}$ T cells. CD3$^{+}$ T cells were subsequently divided into CD4$^{+}$ or CD8$^{+}$ T cells with the expression of CD45RA and CCR7 for naïve and memory phenotype. One representative mouse was shown.

**(B)** Blood of hNSGF6 at 16 weeks after transplant. One representative mouse was shown.

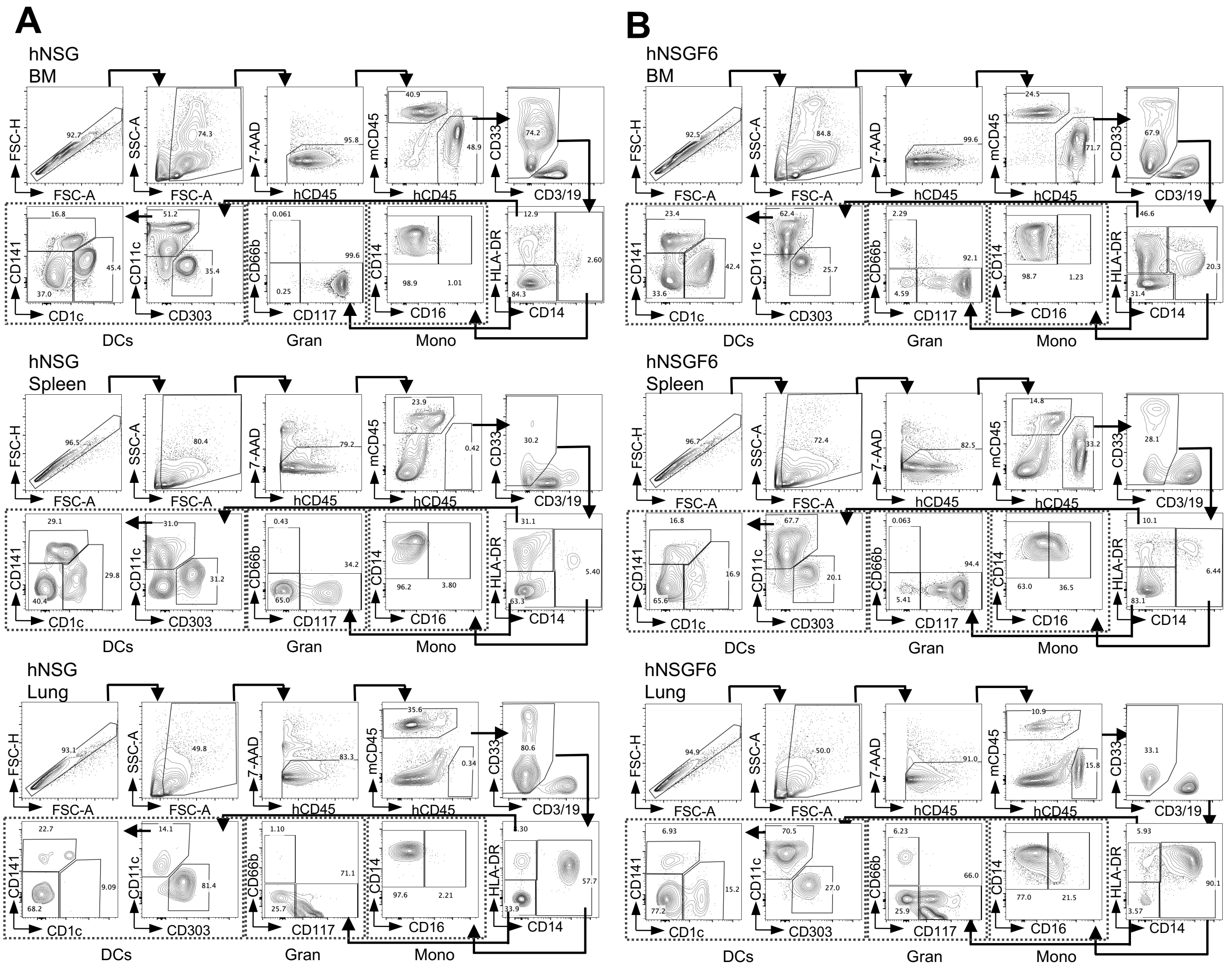


Figure S3. **Evaluation of the human myeloid compartment in the tissues of humanized mice by flow cytometry.** Humanized mice were generated by engrafting mice with 1x10$^5$ bone marrow HPCs. Tissues including bone marrow, spleen and lungs of mice were collected. Single cell suspensions were stained with specific antibodies and analyzed by FACS. Human myeloid cells including DCs, monocytes (Mono) and granulocytes (Gran) were gated in the sequence as shown in the plots.

**(A)** Bone marrow, spleen, and lungs of hNSGF6 at 52 weeks after transplant. One representative mouse was shown.

**(B)** Bone marrow, spleen, and lungs of hNSGF6 at 52 weeks after transplant. One representative mouse was shown.

Table S1. **List of biologicals and reagents used in this study.** See spreadsheet for detail.

Table S2. **List of primers for mouse genotypes.**

| Target | Primer | Sequence | Product (bp) |
|---|---|---|---|
| Flt3 KO | F | GGTACCAGCAGAGTTGGATAGC | 363 (KO) |
| | R | ATCCCTTACACAGAAGCTGGAG | 799 (wt) |
| IL6 KI 5' junction | F1 | CATCTCCTGTGGGACCATTCTTC | 4017 |
| | R1 | AGTGCAGGTTATCTCACTGTGG | |
| IL6 KI 3' junction | F2 | TTGGAACTGAACCCAAGTGTGC | 5163 |
| | R2 | GGCTGTCCTCAGACCCAATC | |
| IL6 KI full-length | F1 | CATCTCCTGTGGGACCATTCTTC | 8155 (KI) |
| | R2 | GGCTGTCCTCAGACCCAATC | 10557 (wt) |
| IL6 KI | F2 | TTGGAACTGAACCCAAGTGTGC | 1475 |
| | R3 | CACTGGCTGGAGTTGGATGC | |
| Donor DNA backbone | F4 | GAAGTTTGTTGCTATGGAAGGGTC | 944 |
| | R4 | AGCGCAACGCAATTAATGTG | |
